# Identification, characterisation and recombinant expression of flavonoid 3’,5’-hydroxylases and cytochrome P450 reductases from *Vaccinium* species

**DOI:** 10.1101/2023.01.09.523147

**Authors:** Kaia Kukk

## Abstract

Flavonoid 3’,5’-hydroxylases (F3′5′Hs) play a key role in biosynthesis of blue coloured anthocyanin complexes in plants. Thus these proteins have potential application in the development of a natural blue coloured food dye using microbial cell factories. However, F3′5′Hs are membrane proteins that require a redox partner, NADPH-cytochrome P450 reductase (CPR). The aim of the research was to identify F3′5′H sequences from *Vaccinium* species plants and express the respective proteins in yeast to test their potential in biotechnological production of precursors of anthocyanins. In this study, novel coding DNA sequences of F3′5′Hs from *Vaccinium myrtillus* and *Vaccinium uliginosum*, and two CPRs from *V. myrtillus* were identified and characterised. The newly obtained proteins and F3′5′H from *Vaccinium corymbosum* and CPR from *Helianthus annuus* were expressed in *Pichia pastoris*. Addition of DMSO into the culture medium increased production of F3′5′Hs and CPRs. A truncated form of *V. corymbosum* F3′5′H, that lacked the predicted first N-terminal alpha helix, expressed at higher level compared to the full-length protein. *Vaccinium* F3′5′Hs were combined with different CPRs and substrates to identify which CPR acts as a redox partner for F3′5′Hs and which substrates are preferred. Unfortunately, only substrates but not the products could be detected, indicating that the recombinant F3′5′Hs were inactive. Therefore, despite progress in protein expression, *P. pastoris* was not a suitable host for producing *Vaccinium* F3′5′Hs.

## INTRODUCTION

Flavonoid 3′,5′-hydroxylases (F3′5′Hs, EC 1.14.14.81) are cytochrome P450 monooxygenases (CYPs) that catalyse hydroxylation of the 3′ and 5′ positions of the B-ring of flavonoids (Figure 1) (Menting, Scopes, and Stevenson 1994; Shimada et al. 1999). Flavonoids are precursors of anthocyanins – the water soluble pigments that give the flowers and fruits red, purple and blue colour. The colour of fruits and flowers is considered to be a crucial factor in attracting seed dispersers and pollinators (Grotewold, 2006). A human being is no difference, being fascinated by colourful flowers and considering bright coloured food as fresh and healthy. The potential health risks of some artificial colourants and health benefits of anthocyanins has been driving the research on employing microbial cell factories for producing anthocyanins to use them as natural food colourants (Koopman et al. 2012; Jones et al. 2017; Appelhagen et al. 2018; Levisson et al. 2018). There is strong interest in obtaining natural blue dye suitable for food industry. Lately, a significant study was published on turning anthocyanins from red cabbage to desired blue coloured anthocyanins using synthetic biology and computational protein design tools (Denish et al. 2021). However, the primary source of anthocyanins was still a plant, red cabbage, making the application seasonable, and the blue complex P2 contained Al^3+^ ions, which raises the question of developing potential aluminium toxicosis after high consumption of food and/or drinks containing such a dye (Igbokwe, Igwenagu, and Igbokwe 2020). There is also the spirulina blue, the phycocyanin from blue-green algae, that is used for giving for example ice cream, drinks and pills the blue colour. However, as phycocyanin is a pigment-protein complex, it is sensitive to elevated temperatures and pH change (Martelli et al. 2014; Chaiklahan, Chirasuwan, and Bunnag 2012). Similarly, the colour of anthocyanins depends on the pH of the environment. Intermolecular copigmentation reactions, acylation and complexing with metal ions and flavones have shown to stabilise anthocyanins (Giusti and Wrolstad 2003; Eiro and Heinonen 2002; Yoshida, Mori, and Kondo 2009).

**Figure 1.**
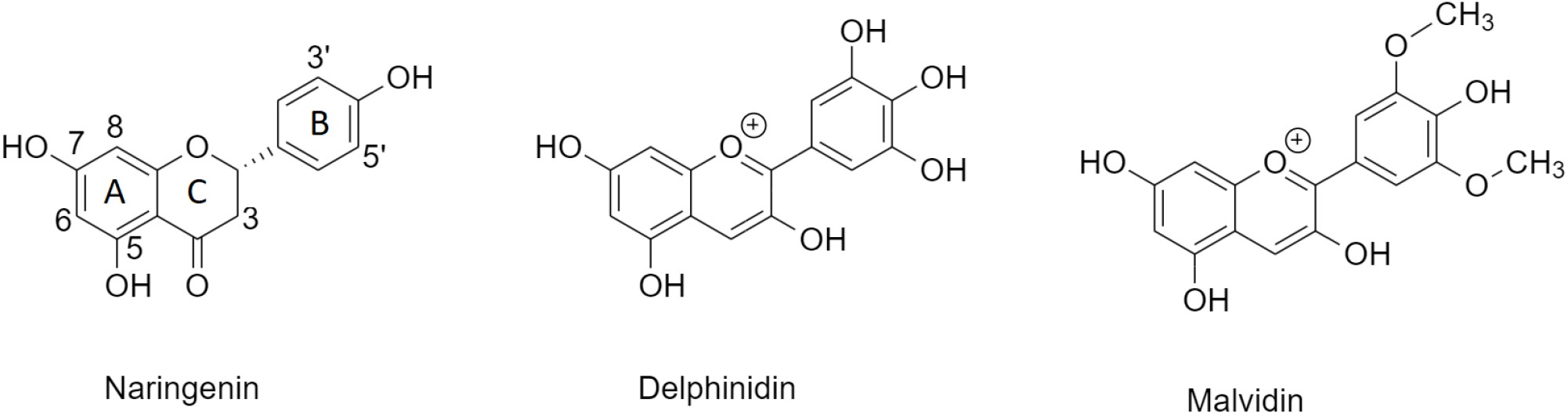
Substrate and downstream products of F3′5′H reaction. An example of a substrate of F3′5′H is naringenin, which is a 3’, 5’ unsubstituted flavanone. Delphinidin is a 3’, 5’ hydroxylated and malvidin a 3’, 5’ methylated anthocyanidin. These anthocyanidins are synthesised from the product of F3′5′H, dihydromyricetin, through several consecutive downstream enzyme reactions.

F3′5′Hs shift the anthocyanin synthesis pathway from the orange and reddish-purple pelargonidin and cyanidin based anthocyanins to purple-blue delphinidin type ones, which makes this enzyme an attractive target for biotechnological applications. The additional hydroxyl groups at 3’ and 5’ positions make way to subsequent modifications (methylation, complexing with metal ions) diversifying the spectrum of anthocyanins. Unfortunately, F3′5′H are complicated objects to study as they are membrane bound proteins tending to inactivate during solubilisation and requiring NADPH-cytochrome P450 reductase (CPR, EC 1.6.2.4) for activity (Menting, Scopes, and Stevenson 1994; Hausjell, Halbwirth, and Spadiut 2018). Published research has used microsomal fractions for characterising protein activity, indicating again difficulties in purifying this protein (Seitz, Ameres, and Forkmann 2007). Studies on F3′5′Hs have focused on the role of these enzymes in blue flower colour development in ornamental plants (Li et al. 2021; Boase et al. 2010), but also gene evolution (Seitz et al. 2006) and regulation of expression (Ma et al. 2021; Karppinen et al. 2021). Recombinant expression of F3′5′Hs has been carried out in tobacco, in *Saccharomyces cerevisiae* and as a fusion protein in *Escherichia coli* (Appelhagen et al. 2018; Seitz, Ameres, and Forkmann 2007; Kaltenbach et al. 1999). Although bilberries (*Vaccinium myrtillus*) and blueberries (*Vaccinium corymbosum*) are rich in malvidin, cyanidin and delphinidin (Müller, Schantz, and Richling 2012) indicating high expression of F3′5′Hs, the F3′5′H genes from *Vaccinium* species are not yet characterised.

The yeast *Pichia pastoris* (*Komagataella phaffii*) is an established system for producing both soluble and membrane bound recombinant proteins as well as value-added chemicals (Cereghino and Cregg 2000; Cregg et al. 2000; Junge et al. 2008; Gao, Jiang, and Lian 2021). The pool of available tools for engineering the expression system is constantly progressing (Z. Yang and Zhang 2018; Fischer and Glieder 2019; Ahmad et al. 2014). There are several selection markers applied to *P. pastoris* such as functional histidinol dehydrogenase gene *HIS4* restoring the ability to grow in histidine deficient media (e.g. used in pPIC9 from Thermo Fisher) and sequences encoding resistance to antibiotics such as G418 (e.g. pPIC9K from Thermo Fisher), zeocin (e.g. pPICZ from Thermo Fisher) and hygromycin B (e.g. pJAH from BioGrammatics). As *HIS4* encoding vectors are rather large (> 8 kb), vectors employing antibiotic resistance that enables direct selection of multi-copy integrants are frequently preferred. In the view of carbon footprint reduction and the amount of electricity required for growing microorganisms, it is worth mentioning that unlike *E. coli, P. pastoris* can be cultivated at room temperature.

In this study obtaining previously uncharacterised coding DNA sequences (CDSs) of CPRs and F3′5′Hs from bilberry (*V. myrtillus*) and bog bilberry (*Vaccinium uliginosum*) is described. Recombinant expression of these proteins and F3′5′H from northern highbush blueberry (*V. corymbosum*) is carried out in *P. pastoris*. Optimisation of culture conditions is carried out and the catalytic activity of the enzymes is assayed.

## MATERIALS AND METHODS

### RNA extraction, complementary DNA synthesis and rapid amplification of cDNA ends for obtaining sequences encoding F3’5’H of *V. myrtillus* and *V. uliginosum*

Spectrum Plant Total RNA Kit (Sigma-Aldrich) was used for extracting RNA from ripe berries of *V. myrtillus* and *V. uliginosum* picked in Western Estonia (58.74772, 24.03531) in July 2019. Complementary DNA (cDNA) was synthesised using RevertAid H Minus First Strand cDNA Synthesis Kit (Thermo Scientific). Sequence fragments were amplified by touchdown PCR (annealing at 72, 65, 58 and 52 °C, number of cycles 5, 5, 5 and 25, respectively) using the following PCR mix: 2 µl cDNA, 1 µl 10 mM dNTPs, 1 µl 100 µM primer F35Hdegsense, 1 µl 100 µM primer F35Hdegantis, 10 µl 5xHF buffer, 0.5 µl Phusion High-Fidelity DNA Polymerase (2 U/µL) (Thermo Scientific) and water to 50 µl. Phusion High-Fidelity DNA Polymerase was used for all PCR experiments of the study. The primers were purchased from Microsynth (Switzerland) or Metabion (Germany). The sequences of all primers are presented in the Supplementary information, Table 1, Primers. *E. coli* DH5ɑ was transformed with PCR products of expected size, purified using GeneJET Gel Extraction and DNA Cleanup Micro Kit (Thermo Scientific) and ligated into pJET1.2 blunt vector (CloneJET PCR Cloning Kit, Thermo Scientific) using T4 DNA ligase (5 U/µl, Thermo Scientific) and propagated. The plasmid DNA was extracted using GeneJET Plasmid Miniprep kit (Thermo Scientific) and subjected to sequencing. In order to perform rapid amplification of cDNA ends (RACE) specific sense, antisense and inner antisense primers were designed according to the obtained sequencing data of fragments. 3’-RACE-ready cDNA was synthesized using RevertAid H Minus Reverse Transcriptase (200 U/µL) (Thermo Scientific) and the primer 3-RACE-OligodT. 3’-RACE-ready cDNA and specific sense primer in combination with the primer 3-RACE-spec were used to amplify 3’ sequences of F3′5′H (conventional PCR according to manufacturer’s instructions). FirstChoice RLM-RACE Kit (Invitrogen), gene specific antisense and inner antisense primers were used to obtain 5’ sequences. The procedure is described in more detail in the protocol published on protocols.io (Kukk 2021).

### Amplification of potential CPR sequences from *V. myrtillus* and *Helianthus annuus* and synthesis of *V. corymbosum* F3′5′H

RNA was extracted from leaves of *H. annuus* picked from plants grown in outdoor conditions and from berries of *V. myrtillus* using Spectrum Plant Total RNA Kit (Sigma-Aldrich). cDNA was synthesized using RevertAid H Minus First Strand cDNA Synthesis Kit (Thermo Scientific) and oligo(dT)18 primer. The predicted sequence of *Helianthus annuus* CPR (*Ha*CPR, XM_022125659.2) was used as a template for designing specific 5’ and 3’ primers HaCPR5p and HaCPR3p. The Blast+ tool on Vaccinium.org page was used to identify potential CPR sequences from the genomic data of *V. myrtillus* (Wu et al. 2022). Specific primer pairs were designed for two of protein coding sequences exhibiting expected length and the highest similarity to the predicted *Ha*CPR: *Vm*CPR15p and *Vm*CPR13p, and *Vm*CPR25p and *Vm*CPR23p. PCR was carried out according to manufacturer’s instructions using 2 µl cDNA per 50 µl reaction. The PCR products of correct size were ligated into pJET1.2 blunt, used for transforming *E. coli* DH5ɑ, propagated and sequenced.

The sequence encoding *V. corymbosum* F3′5′H (*Vc*F3′5′H, GenBank accession number MH321464.1) was used as a template to order a synthetic sequence codon-optimised for yeast expression (BioCat GmbH, GenBank accession number ON695863).

### Construction of a *P. pastoris* expression vector encoding hygromycin B resistance marker

A 4942 bp expression vector named pAOXHygR (Supplementary information, file pAOXHygR.txt) encoding alcohol oxidase 1 (AOX1) promoter, multiple cloning site (MCS) followed by *myc* epitope and a 6xHis tag, hygromycin B resistance marker, origin of replication and ampicillin resistance marker was constructed by Golden Gate assembly. Vectors pPICZ A (Invitrogen), pUDP082 (Addgene plasmid #103875) (Juergens et al. 2018) and pJET1.2, and the primer pairs prAOX1MCSHis6TT FW and prAOX1MCSHis6TT REV, pHygRt FWD and pHygRt REV, and ori pAmpRt FWD and ori pAmpRt REV were used to amplify the fragments to be assembled. The BsaI restriction site in the ampicillin resistance gene was removed by introducing a silent mutation (GGG to GGA). Golden Gate assembly was carried out by incubating PCR products purified from agarose gel with BsaI, T4 DNA ligase and the buffer of the T4 DNA ligase. The restrictases used in the study were from Thermo Scientific. The *E. coli* DH5α transformants were selected on LB agar plates containing 75 µg/ml ampicillin. The vector construction procedure is described in more detail in a protocol published on protocols.io (Kukk 2022a).

### Plasmid construction for expressing F3′5′Hs and CPRs in *P. pastoris*

The sequences of *V. myrtillus* and *V. uliginosum* F3′5′Hs (*Vm*F3′5′H and *Vu*F3′5′H) were fused with the Kozak consensus sequences and the N-terminal hexahistidine (6xHis) tag by PCR using primers *Vm*F35HEcoRIKozNtH6 and VmF35HCtermEcoRI, and VuF35HEcoRIKozNtH6 and VuF35HCtermEcoRI, respectively. Ligation into the EcoRI restriction site of pPICZ A vector (Invitrogen, Thermo Fisher Scientific) followed. The synthetic sequence of *Vc*F3′5′H contained an N-terminal 6xHis tag followed by the NcoI restriction site. For inserting *Vc*F3′5′H fused with Kozak consensus sequence and the tag into the ApaI site of pPICZ A, PCR was carried out with the primers *Vc*F35HApaIKozakNtH6 and *Vc*F35HCtermApaI. C-terminally affinity tagged *Vaccinium* F3′5′Hs were generated by removing the stop codon and inserting the sequences into the EcoRI or ApaI sites of the pPICZ A vector in frame with the *myc* epitope and the C-terminal 6xHis tag. The following primer pairs were used in PCR: *Vm*F35HEcoRIKozwoH6 and *Vm*F35HCtermnostopEcoRI, *Vu*F35HEcoRIKozwoH6 and *Vu*F35HCtermnostopEcoRI, and *Vc*F35HApaIKozwoH6 and *Vc*F35HCtermnostopApaI. In the case of *Vm*F3′5′H and *Vu*F3′5′H, the vector sequence between the EcoRI site and *myc* epitope had been removed prior ligation, so that there was no extra sequence except the restriction site between the coding sequences of all three F3′5′Hs and the *myc* epitope of the vector.

N-terminally truncated sequences of *Vaccinium* F3′5′Hs (lacking residues Ala2-Gln30) were generated by PCR using the following primer pairs: *Vm*F35HApaIwoTMH and *Vm*F35HApaInostoprev, *Vu*F35HApaIwoTMH and *Vu*F35HApaInostoprev, and *Vc*F35HApaIwoTMH and *Vc*F35HCtermnostopApaI. As before, Kozak consensus sequence was fused to 5’ end of the sequence and the stop codon was removed for affinity tagging. The sequences were ligated into the ApaI site of pPICZ A.

The VmCPR sequences were ligated into the ApaI site of the newly generated pAOXHygR vector. HaCPR was ligated into NotI site of pAOXHygR. The primer pairs VmCPR1ApaIfw and VmCPR1ApaIrev, VmCPR2ApaIfw and VmCPR2ApaIrev, and HaCPRNotIfw and HaCPRNotIrev were used to introduce the restriction sites and the Kozak consensus sequence, and remove the stop codon for fusing the sequences with the C-terminal *myc* epitope and a 6xHis tag from the vector.

For expressing N-terminally 6xHis tagged CPRs, the following primer pairs were used: VmCPR1ApaINtH6fw and VmCPR1ApaIstoprev, VmCPR2ApaINtH6fw and VmCPR2ApaIstoprev, and HaCPRNotINtH6fw and HaCPRNotIstoprev. The PCR products were digested with ApaI or NotI and ligated into the same sites of pAOXHygR vector.

*E. coli* DH5α was transformed with the plasmids and selection was carried out on low salt LB plates (10 g tryptone, 5 g NaCl, 5 g yeast extract per litre, pH 7.5) containing 25 µg/ml zeocin (Invitrogen, Thermo Fisher Scientific) when pPICZ A had been used or conventional LB plates (10 g tryptone, 10 g NaCl, 5 g yeast extract per litre) containing 75 µg/ml ampicillin for pAOXHygR plasmids.

### Yeast transformation

Prior yeast transformation, pAOXHygR expression vectors containing CPR sequences were linearised with MssI (PmeI) and pPICZ A vectors encoding F3’5’Hs with BglII. 2-2.5 μg of linear DNA was used for transforming *P. pastoris* GS115H [GS115 with *HIS4* phenotype obtained by transforming GS115 (his4, Thermo Scientific) with pPIC3.5 (Thermo Scientific) linearised with StuI]. Sufficient amount of plasmid DNA was generally extracted from one or two 1.5 ml *E. coli* DH5ɑ cultures. Electroporation was principally carried out as before, incorporating pretreatment of the cells with dithiothreitol (Kukk and Samel 2016). However, in this study the cells were plated onto the YPDS plates (1% yeast extract, 2% peptone, 2% dextrose, 2% agar, 1 M sorbitol) containing 100 or 500 µg/ml zeocin or YPD (YPDS without sorbitol) containing 200 µg/ml hygromycin B (Invitrogen, Thermo Fisher Scientific) already after 2 hours of incubation in 1 M sorbitol. The plates were incubated at 29 °C for two to three days or at room temperature for three to four days. Then, colonies were streaked onto fresh YPD plates containing 100, 500 or 1000 µg/ml zeocin or 200 µg/ml hygromycin B and incubated for 24 hours. Three colonies per construct were picked to generate glycerol stocks and test protein expression. Colony PCR with gene specific primers was carried out to confirm integration of the gene of interest.

### Protein expression, optimisation and analysis

Preliminary expression experiments were carried out in autoinduction medium (Lee et al. 2017). A single colony was used to inoculate 10 ml of the medium containing 1% yeast extract, 2% peptone, 100 mM potassium-phosphate buffer (pH 6), 8 × 10^−5^% biotin, 2.68% yeast nitrogen base with ammonium sulfate and without amino acids (YNB), 0.4% glycerol and 0.5% methanol. The culture was incubated at 25 °C for 48 hours. Addition of several supplements (0.004% histidine, 10 µM hemin, 2.5% dimethyl sulfoxide (DMSO)) was tested to improve the expression of full-length *Vaccinium* F3′5′Hs. For disrupting yeast cells, approximately 130 mg (wet weight) of cells were resuspended in 0.5 ml of lysis buffer containing 50 mM Tris-HCl (pH 8.0), 5 mM EDTA and 1 mM PMSF. Five cycles of sonication for 10 s at a power setting of 5 (Torbeo Ultrasonic Cell Disruptor) followed. Cell lysates of induced yeast cells were analysed by SDS-PAGE and Western analysis essentially as before (Kukk and Samel 2016), except PVDF membrane (0.45 μm, Immobilion-P, Millipore) was used. PageRuler prestained protein ladder (Thermo Scientific) was used for molecular weight estimation. Protein structure was predicted using Alphafold v2.1.0 (Jumper et al. 2021). For linking protein expression with mRNA levels, RNA was extracted from induced yeast cells using Spectrum Plant Total RNA Kit (Sigma-Aldrich) and cDNA was synthesised using RevertAid H Minus First Strand cDNA Synthesis Kit (Thermo Scientific). Then, PCR was run with 5′ and 3′ AOX1 sequencing primers. *N-*glycosylation was confirmed by PNGase F treatment (Kukk 2022b).

### Activity assay

50 μl of the lysates of yeast cells expressing F3’5’H and CPR were pipetted into a glass vial with a magnetic stir bar. 10 μM substrate (naringenin, eriodictyol, dihydrokaempferol, dihydroquercetin, kaempferol) dissolved in ethanol and 1 mM NADPH was added. The final volume was 500 μl and the reaction was carried out in 50 mM Tris-HCl, pH 8. The reaction mix was incubated with stirring at room temperature for 40 min. The products of the enzyme reaction were extracted twice with 500 μl of ethyl acetate, evaporated and dissolved in 50 μl of ethanol.

Chromatographic separation was carried out using Agilent Infinity 1290 HPLC system, Agilent Zorbax Extend C18 column (2.1 × 50 mm, 1.8 μm) and a binary mobile phase system. Mobile phase A was water containing 0.1% formic acid (v/v) and mobile phase B was methanol. The elution program was as follows: 20 % B, 0 - 3 min; 50 % B, 3 - 8 min; 90 % B, 8 - 11 min; 10 % B, 11 - 16 min. The flow rate was 0.2 ml/min and the sample injection volume 1 μl. The oven temperature was 45 °C. Agilent Technologies 6540 UHD Accurate-Mass Q-TOF LC/MS spectrometer and electrospray ionization was used for identifying the substrates and products of the enzyme reactions.

## RESULTS AND DISCUSSION

### RNA extraction from berries of *V. myrtillus* and *V. uliginosum*

Obtaining novel CDSs from plants that do not have genomic data available requires RNA extraction and complementary DNA (cDNA) synthesis. At the beginning of this study, the genome of *V. myrtillus* (Wu et al. 2022) was not available yet and the genome of *Vaccinium uliginosum* is still not. Extraction of RNA from anthocyanin and polysaccharide rich plant material may be complicated. At first I tried the protocols developed for *Arabidopsis thaliana* (Oñate-Sánchez and Vicente-Carbajosa 2008) which worked fine for extracting RNA from leaves of, for example, sunflower, but not for bilberries. Application of protocol 2 from the same article resulted in RNA that, according to agarose gel electrophoresis, had high quality, however the low ratio of absorbance at 260 and 280 nm indicated contamination with phenol. Finally, switching to a commercial kit Spectrum Plant Total RNA Kit from Sigma-Aldrich gave approximately 2.5 μg of RNA from 100 mg of berry powder. The yield may be considered modest, however, it was sufficient for downstream experiments. The same kit was also suitable for extracting RNA from yeast, the yield was however more than forty times higher than from berries.

### Characterisation of novel F3’5’Hs and CPRs from *Vaccinium* species

The degenerate primers (Supplementary information, Table 1, Primers) generated for the study were successfully used for amplifying fragments of potential F3′5′Hs from *Vaccinium* species which led to isolation of full-length CDSs of F3′5′Hs from the berries of *V. myrtillus* and *V. uliginosum*. I would like to make a remark that these degenerate primers may not be universal, as they were not suitable for obtaining F3′5′H fragments from blue flowers, such as *Centaurea cyanus* and *Cichorium intybus*. The obtained sequences were deposited in GenBank with the following accession numbers: OK533469.1 (*Vm*F3′5′H), OK533468.1 (*Vu*F3′5′H) and MW837166.1 (partial sequence of another isoform of *Vm*F3′5′H). The full-length CDSs of *Vm*F3′5′H and *Vu*F3′5′H consist of 1539 basepairs and they share ~95% sequence identity with F3′5′H from *V. corymbosum* (MH321464.1), ~84% sequence identity with F3′5′Hs from *Rhododendron* species and ~76% sequence identity with F3′5′Hs from *Vitis* species. The amino acid sequences of F3′5′Hs from the three *Vaccinium* species are 95-97% identical, whereas the sequences of *Vu*F3′5′H and *Vc*F3′5′H have slightly higher sequence identity compared to other combinations (Supplementary information, file Alignment of *Vaccinium* F3′5′H proteins). The partial sequence of the other isoform of *Vm*F3′5′H (MW837166.1) shares 85% nucleotide sequence identity with the “first” isoform isolated with the full-length CDS (OK533469.1). The predicted structure models of *Vaccinium* F3′5′Hs (Supplementary information, files Vc/Vm/VuF35H.pdb) are virtually overlapping and show a structure with a protruding alpha helix possibly acting as a membrane anchor (Figure 2, A). The overall structure of *Vaccinium* F3′5′Hs is similar to the structure of other CYPs (Guengerich, Waterman, and Egli 2016).

**Figure 2.**
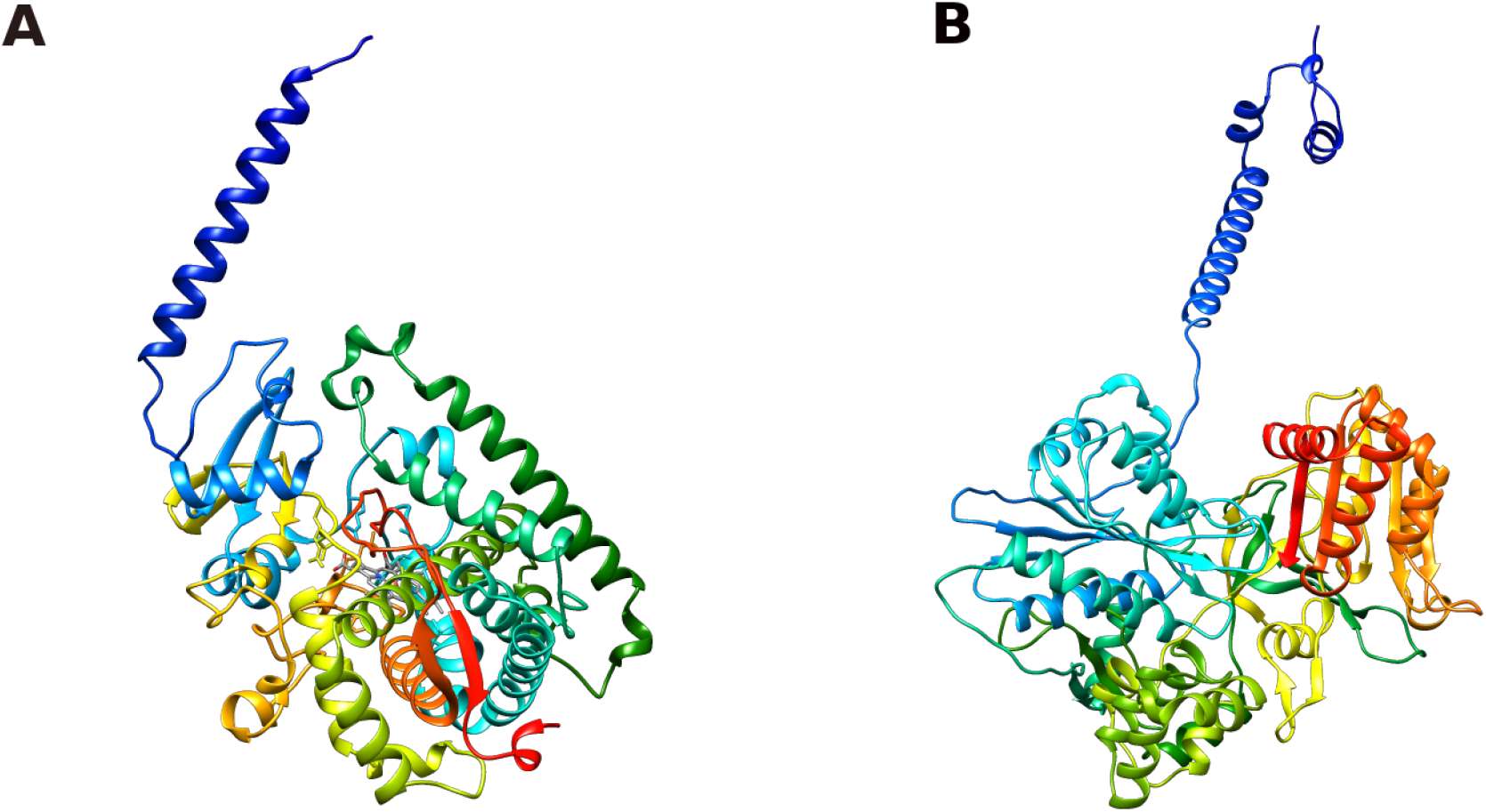
Predicted tertiary structures of A) *Vc*F3’5’H and B) *Vm*CPR2. Rainbow colouring is used whereas the N-terminus is marked with blue and C-terminus with red. The model of *Vc*F3’5’H is positioned likewise CYP 21A2 in figure by Guengerich et al. (Guengerich, Waterman, and Egli 2016) to demonstrate the similarity of the structure of *Vc*F3’5’H, a plant CYP, to a mammalian counterpart, CYP 21A2. The model prediction of *Vc*F3’5’H and *Vm*CPR2 was carried out with Alphafold v2.1.0 (Jumper et al. 2021) and the image was created using UCSF Chimera (Pettersen et al. 2004).

The homology search against the genome of *V. myrtillus* revealed two potential CPR sequences that were used for designing specific primers. After amplification from cDNA of *V. myrtillus* the obtained potential CPR sequences were sequenced and compared with the genomic data of *V. myrtillus*. One of the isoforms (*Vm*CPR2, GenBank accession number OP729290) was identical with the sequence from the published genome, the other (*Vm*CPR1, GenBank accession number OP729289) had the following nucleotide substitutions: T666G causing His222Gln and the silent mutations T1845C and A1893G. The CDSs of predicted *Vm*CPR1 and *Vm*CPR2 consist of 2127 and 2109 bp. Both isoforms share ~80% sequence identity with other plant CPRs. The CPR sequence from sunflower, *H. annuus*, used in the study in encoded by 2124 bp. The amino acid sequence identity between *Ha*CPR and *Vm*CPRs is 77-78%. The predicted tertiary structure of *Ha*CPR (Uniprot entry A0A251TTX6) is similar to the predicted structures of *Vm*CPR1 and *Vm*CPR2 (Figure 2, B, Supplementary information, files VmCPR1.pdb and VmCPR2.pdb), and similarly to F3’5’Hs, CPRs have a protruding alpha helical structural motif that is associated with membrane binding.

### Expression of *Vaccinium* F3’5’Hs and CPRs in *P. pastoris*

*P. pastoris* was transformed with both N- or C-terminally affinity-tagged *Vaccinium* F3’5’Hs. Subsequent expression experiment revealed that *Vaccinium* F3’5’Hs fused with the N-terminal 6xHis tag were barely detectable. F3’5’Hs are located in the microsomal fraction and likely contain an N-terminal signal-anchor that is not cleaved similarly to other CYPs (Szczesna-Skorupa and Kemper 2011). In addition, prediction of a cleavable signal peptide using SignalP v 6.0 (Teufel et al. 2022) resulted no hits. Therefore, the inability to detect *Vaccinium* F3’5’Hs was not likely due to N-terminal truncations removing the affinity tag. It may be speculated that the N-terminal 6xHis tag interferes with membrane insertion of F3’5’Hs and causes subsequent misfolding and degradation of the protein.

*Vaccinium* F3’5’Hs fused with C-terminal *myc* epitope and 6xHis tag from the vector were produced at different levels. The expression of *Vm*F3’5’H was very low, whereas *Vc*F3’5’H and *Vu*F3’5’H exhibited modest expression. Although the sequence of *Vc*F3’5’H was codon optimised for yeast, *Vu*F3’5’H encoded by its native sequence was produced at similar level (Figure 4, A). Therefore, at first sight, codon optimisation seemed not to be critical for expressing plant enzymes in *P. pastoris*. Human proteins with only slightly higher codon adaptation indexes (CAI) have been expessed in *P. pastoris* at decent levels (CAI 0.63-0.64 for *Vaccinium* F3’5’Hs vs. for example 0.69 for pancreatic lipase (Villo et al. 2020) and 0.70 for prostaglandin H synthase (Kukk and Samel 2016)). In order to clarify what is behind poor expression of *Vm*F3’5’H, I compared the mRNA levels of F3’5’Hs by running PCR with AOX1 sequencing primers and cDNA extracted from yeast cells that had been induced to express these F3’5’Hs. The amount of the PCR product correlated with the protein production, which indicated that mRNA stability may be the key factor and sequence optimisation may carry more value than considered in the first place. Although the calculated molecular weights of *Vc*F3’5’H and *Vu*F3’5’H differ only by 0.3 kDa, according to SDS-PAGE *Vu*F3’5’H appears to have ~4 kDa smaller molecular weight than *Vc*F3’5’H. Therefore, the PCR products obtained for evaluating mRNA levels were also subjected to sequencing, which re-confirmed that the sequence of *Vu*F3’5’H was full-length.

In order to increase the yields of F3’5’Hs, the effect of several supplements (histidine, hemin, DMSO) and expression conditions on the production was evaluated. Compared to the conventional BMGY/BMMY media, the autoinduction medium used in the study contained double amount of YNB and biotin, 2.5 times less glycerol and methanol already from the start (Lee et al. 2017). Based on band intensities on immunoblotted membrane, expression in autoinduction medium supplemented with 2.5% of DMSO versus non-supplemented autoinduction medium yielded three times more *Vc*F3’5’H and *Vu*F3’5’H, whereas supplementing the conventional BMMY medium with DMSO did not have such an effect. Higher expression level was also not due to increased amount of YNB, as parallel experiments with media containing 1xYNB and 2xYNB showed no significant difference. Therefore, the mixed feeding strategy with glycerol and methanol was necessary for producing these proteins, probably due to slowing down the expression of the recombinant proteins thereby not overwhelming the (re-)folding machinery of the yeast. Positive effect of DMSO on expression level of membrane proteins has been shown before in yeast as well as mammalian cell lines (André et al. 2006; Shukla, Reinhart, and Michel 2006). Addition of hemin and histidine did not have noticeable effect on protein production.

The predicted structure of F3’5’Hs shows an N-terminal alpha helix that protrudes from the protein (Figure 2, A). I decided to test whether removing this helix (residues Ala2-Gln30) affects production of recombinant *Vaccinium* F3’5’Hs and membrane binding. Surprisingly, truncated *Vc*F3’5’H expressed at noticeably higher level than the full-length enzyme, and in conventional BMGY/BMMY media. Inclusion of DMSO into the medium did not increase the expression of truncated *Vc*F3’5’H noticeably indicating that membrane fluidity may not be so important for folding of this protein. However, it still mostly remained in insoluble fraction and poorly solubilised in the presence of detergents. Unexpectedly, truncated *Vm*F3’5’H and *Vu*F3’5’H were not produced at detectable levels. It is difficult to bring out the reason why N-terminal truncation increased the production of *Vc*F3’5’H but eliminated *Vm*F3’5’H and *Vu*F3’5’H. It could be speculated that the truncation made the mRNA of *Vm*F3’5’H and *Vu*F3’5’H more unstable, but in the optimised sequence of *Vc*F3’5’H such instability element(s) had been removed. Positive effect of truncation on expression of *Vc*F3’5’H may indicate that membrane insertion is the bottleneck in *Vaccinium* F3’5’H production in *P. pastoris*. Expression experiments with optimised sequences of truncated *Vm*F3’5’H and *Vu*F3’5’H would give answers to these questions, but at the final stage of the study these were not financially feasible.

A frequently used strategy when CYPs are expressed recombinantly is co-expression with CPRs (Hausjell, Halbwirth, and Spadiut 2018) or even as fusion proteins with CPR (Kaltenbach et al. 1999). For carrying out co-expression of CPRs and F3’5’Hs I decided to construct a yeast expression vector pAOXHygR (Figure 3) encoding hygromycin B selectable marker. Utilisation of hygromycin B resistance for selecting transformants of *P. pastoris* has been described before (J. Yang et al. 2014), but application is not so common yet.

**Figure 3.**
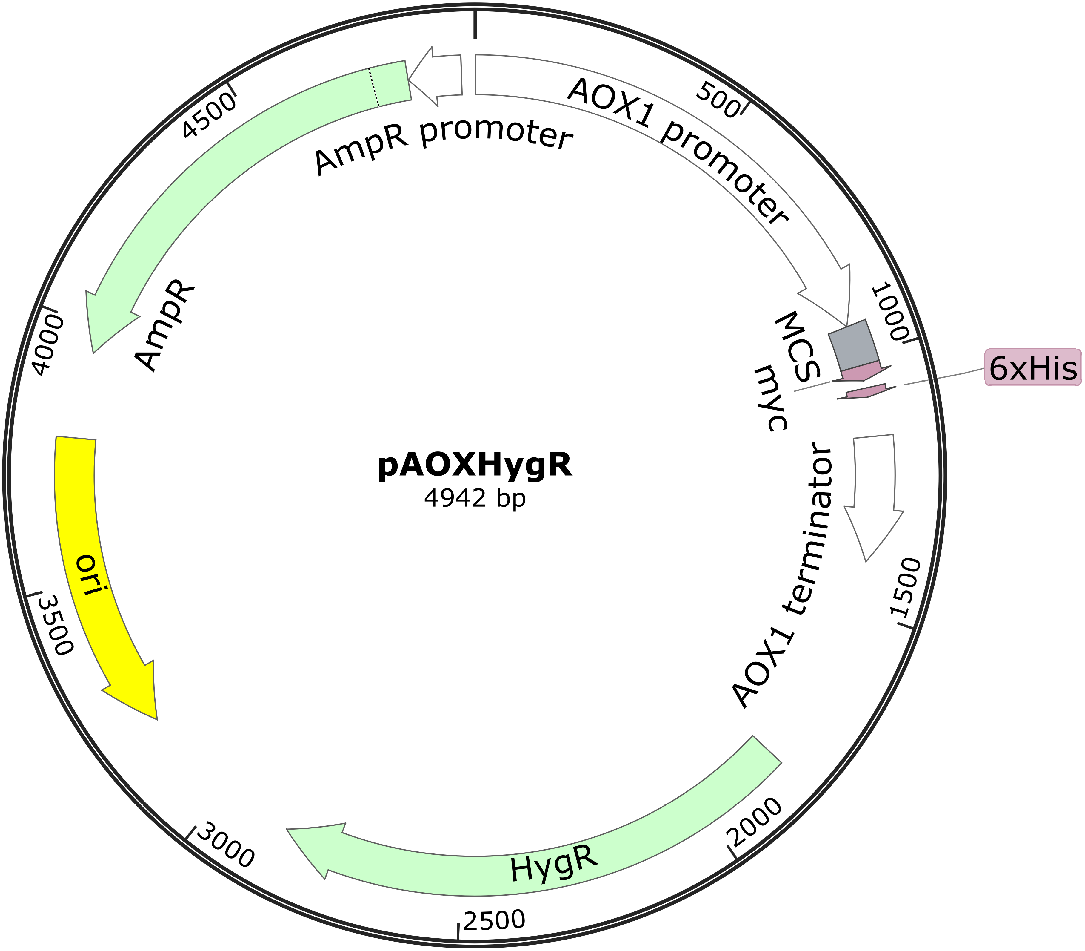
Map of pAOXHygR vector showing the main features of the vector: AOX1 promoter, MCS, *myc* epitope, 6xHis tag, AOX1 terminator, hygromycin B resistance marker, origin of replication and ampicillin resistance marker. The image was created with SnapGene Viewer 6.1.2.

CPRs have a conserved C-terminal tryptophan residue that is proposed to participate in NADPH binding (Zhang et al. 2022). Therefore, in addition to C-terminally affinity tagged CPRs, N-terminally affinity tagged forms were also expressed to test whether C-terminal tag affects the functionality of the CPRs. Both CPRs from *V. myrtillus* (*Vm*CPR1 and *Vm*CPR2) and the CPR from *H. annuus* (*Ha*CPR) expressed at more or less similar levels independent of the tag location. Western analysis of the strains expressing CPRs fused with the N-terminal 6xHis tag is shown of Figure 4, B. Similarly to F3’5’Hs, addition of DMSO into the culture media increased expression of CPRs. *Vm*CPR1 and *Ha*CPR sequences contain two *N*-linked glycosylation consensus sequences and these proteins are *N*-glycosylated when expressed in *P. pastoris*. Multiple glycoforms of *Vm*CPR1 are visible on Figure 4, B. *N-*glycosylation of *Ha*CPR was also confirmed by PNGase treatment, which reduced the molecular weight of the protein. Western analysis also reveals that expression levels of the recombinant proteins vary noticeably when strains are selected on plates containing 200 μg/ml hygromycin B. Higher concentrations of the antibiotic were not used in this study.

**Figure 4.**
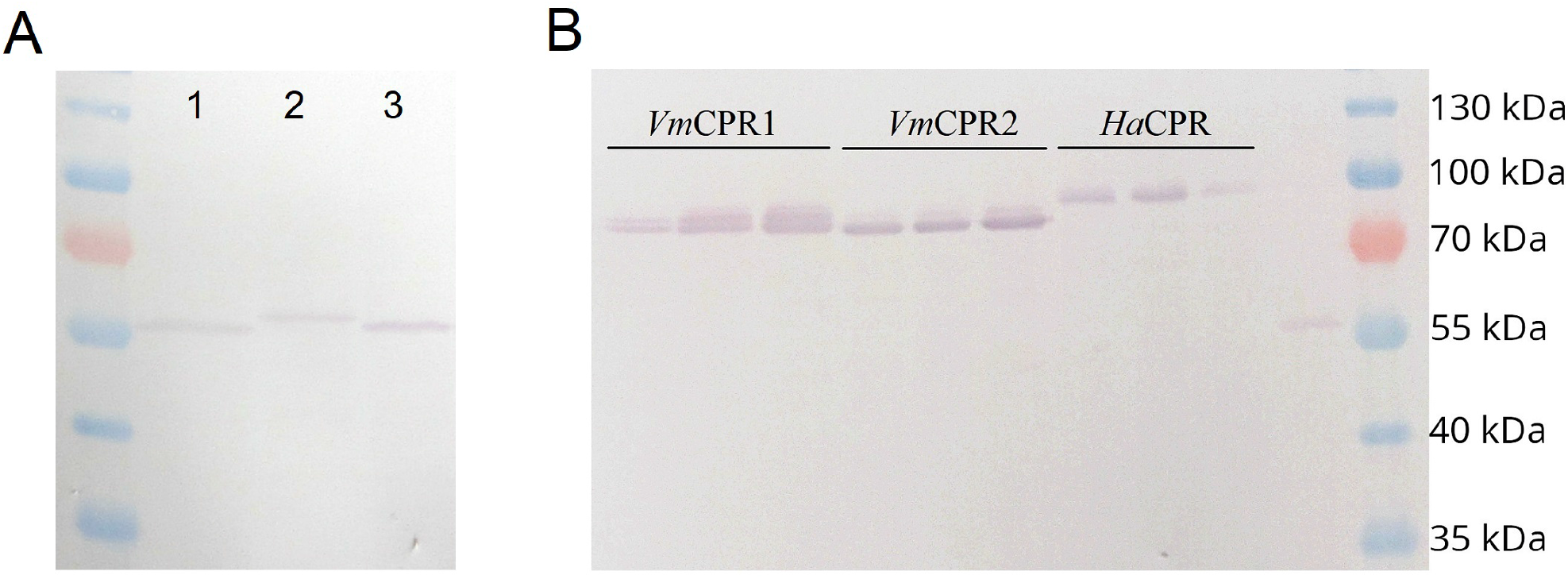
Western analysis of microsomes of *P. pastoris* cells expressing A) C-terminally affinity tagged 1) *Vu*F3’5’H, 2) *Vc*F3’5’H and 3) truncated *Vc*F3’5’H at optimised conditions, and B) N-terminally affinity tagged CPRs from *V. myrtillus* and *H. annuus*. Three transformed strains selected on 200 μg/ml hygromycin B are shown. The calculated molecular weight of F3’5’Hs, truncated *Vc*F3’5’H and CPRs are ~60, 56 and ~79 kDa, respectively. PageRuler prestained protein ladder is on the right.

Transformants that had integrated both F3’5’H and CPR sequences exhibited resistance to zeocin and hygromycin B. The expression of the proteins was induced with methanol. According to Western analysis co-expression strains produced less F3’5’H and CPR than the strains that expressed one protein at a time. Thus, the yeast was producing recombinant proteins at its upper limit and co-expression brought no benefits. Therefore, cell lysates of strains expressing only F3’5’H or CPR were combined for activity measurements.

### Activity assays of *Vc*F3’5’H and *Vu*F3’5’H

The activity of *Vaccinium* F3’5’H was determined by incubating the yeast lysates of the cells expressing F3’5’H and CPR (N- or C-terminally affinity tagged), NADPH and a substrate. Naringenin, eriodictyol, dihydrokaempferol, dihydroquercetin and kaempferol were tested as potential substrates. The formation of products was monitored by HPLC-MS. Unfortunately, despite the efforts made for enhancing the expression of these proteins in a foreign host, I was unable to detect formation of products of the hydroxylation reaction. Elution profiles showing separation of standard compounds naringenin and eriodictyol, and flavonoids extracted from reaction mix containing naringenin and yeast lysates (Supplementary information, HPLC chromatograms) confirm that the method was suitable for separating flavonoids hydroxylated at different number of positions. In addition, mass spectrometry data confirmed that the substrate had not been hydroxylated.

## CONCLUSION

In this study, sequences encoding potential F3’5’Hs from *V. myrtillus* and *V. uliginosum*, and CPRs from *V. myrtillus* were identified and subjected to recombinant expression in *P. pastoris*. F3’5’Hs were produced at poor levels, however optimisation of expression conditions improved the yields noticeably. Unfortunately, F3’5’Hs exhibited no activity towards naringenin and other substrates. Hausjell et al. conclude in their review that compared to *P. pastoris*, higher yields of CYPs have been achieved in *S. cerevisiae* (Hausjell, Halbwirth, and Spadiut 2018). This study confirms that view. The results did not satisfy the expectations and raised question about the probability of catching a F3’5’H gene that could be easily produced in a microbial cell factory for synthesising precursors of anthocyanins. Computational tools and thorough knowledge of the active site structure of F3’5’Hs could lead to designing of an optimised protein exhibiting expected catalytic activity and good expression levels, however this a topic of another study.

## Supporting information

Suppl protein structure 1

Suppl protein structure 2

Suppl protein structure 3

Suppl protein structure 4

Suppl protein structure 5

Suppl prot alignment

Suppl HPLC

Suppl Vector sequence

## ABBREVIATIONS

AOX1: alcohol oxidase 1
cDNA: complementary DNA
CDS: coding DNA sequence
CPR: NADPH-cytochrome P450 reductase
CYP: cytochrome P450 monooxygenase
DMSO: dimethyl sulfoxide
F3′5′H: flavonoid 3′,5′-hydroxylase
Ha: *Helianthus annuus*
MCS: multiple cloning site
RACE: rapid amplification of cDNA ends
*Vc*: *Vaccinium corymbosum*
*Vm*: *Vaccinium myrtillus*
*Vu*: *Vaccinium uliginosum*.

## DATA ACCESSIBILITY

The primer sequences, HPLC chromatograms and the sequence of pAOXHygR vector are provided in Supplementary information as pdf and txt files. The predicted tertiary structures of *V. myrtillus* and *V. uliginosum* F3’5’Hs and *V. myrtillus* CPRs are available in Supplementary information as pdb files.

## AUTHOR CONTRIBUTIONS

KK carried out design of the study, experimental work, data analysis and wrote the manuscript.

## ACKNOWLEDGMENTS

This work was supported by the European Regional Development Fund (ERDF) project Nr.1.1.1.2/VIAA/2/18/286 “Optimization of novel plant derived enzyme expression in microorganisms for biotechnological application. I would like to thank Ivar Järving for instructing me in HPLC-MS.

