## Supplementary material for "Identification, characterisation and recombinant expression of flavonoid 3’,5’-hydroxylases and cytochrome P450 reductases from *Vaccinium* species": Suppl prot alignment

CLUSTAL O(1.2.4) multiple sequence alignment of *Vaccinium* F3’5’Hs
Alignment created on https://www.ebi.ac.uk/Tools/msa/clustalo/
Madeira F, Pearce M, Tivey ARN, et al. Search and sequence analysis tools services from EMBL-EBI in 2022. Nucleic Acids Research. 2022 Apr:gkac240. DOI: 10.1093/nar/gkac240. PMID: 35412617; PMCID: PMC9252731.

```
VmF3'5'H      MALDIILLREIAAATVVFLLTRFFLGSLLQLLKPASKLPPGPKGWPVVGALPLLGTMPHV      60
VuF3'5'H      MALDIMLFREIAAATVVFLLTRFFLGSILQLLKPTCKLPPGPKGWPVVGALPLLGTMPHV      60
VcF3'5'H      MAPDIMLFREIAAATVVFLLTRLFLGSILQLLKPTCKLPPGPKGWPVVGALPLLGTMPHV      60
              ** **:*:**************:****:******:.************************

VmF3'5'H      ALAKMAKKYGPIVYLKMGTCGMVVASTPESARAFLKTLDMNFSNRPPNAGATLLAYNSQD      120
VuF3'5'H      ALAKMAKKYGPIVYLKMGTCGMVVASTPESARAFLKTLDMNFSNRPPNAGATHLAYNSQD      120
VcF3'5'H      ALAKMAKKYGPIVYLKMGTCGMVVASTPESARAFLKTLDMNFSNRPPNAGATHLAYNSQD      120
              **************************************************** *******

VmF3'5'H      MVFADYSPKWTLLRKLCNVHMLGGKALDDSAHIRESELGHMLRAMVESSQRAEPVVISEM      180
VuF3'5'H      MVFADYGPKWTLLRKLCNVHMLGGKALEDSAHIRESELGHMLRDMVELSQRAEPVVISEM      180
VcF3'5'H      MVFADYGPKWTLLRKLCNVHMLGAKALDDSAHIRKSELGHMLRAMVESSQRAEPVVISEM      180
              ******.****************.***:******:******** *** ************

VmF3'5'H      MTYAMANMIGQVILKRRVFVQKGLESNEFKDMVVELMTSAGLFNVGDFIPAIAWMDLQGI      240
VuF3'5'H      MTYAMANMIGQVILKRRVFVQKGSESNEFKDMVVELMTSAGLFNVGDFIPSVAWMDLQGI      240
VcF3'5'H      MTYAMANMIGQVILKRRVFVQKGSESNEFKDMVVELMTSAGLFNVGDFIPSVAWMDLQGI      240
              *********************** **************************::********

VmF3'5'H      EGGMKRMHKKWDDLITRMVKEHAESALERKGNPDFLDVLMANKENSQGTGSLSLTNIKAL      300
VuF3'5'H      EGGMKRMHSKWDYLITRMVKEHSESAHERKGNPDFLDVLMANKENSQGTGSLSLTNIKAL      300
VcF3'5'H      EGGMKRMHSKWDDLITRMVKGHSESAHERKGNPDFLDVLMANKENSQGIGSLSLTNIKAL      300
              ********.*** ******* *:*** ********************* ***********

VmF3'5'H      LLNLFTAGTDTSSSVIEWALAEMVLNPNILKRTQQEMDQVIGRNRRLQESDISKLPYLQA      360
VuF3'5'H      LLNLFTAGTDTSSSVIEWALSEMLLNPDILKRAQQEMDQVIGRNRRLKESDIPKLPYLQA      360
VcF3'5'H      LLNLFTAGTDTSSSIIEWALAEMVLNPDILKRAQQEMDQVIGRNRRLQESDIPKLPYLQA      360
              **************:*****:**:***:****:**************:**** *******

VmF3'5'H      LCKEAFRLHPSTPLNLPRISSEACEVNGYYVPKNTRLSVNIWAIGRDPDVWENPLEFNPE      420
VuF3'5'H      LCKEAFRLHPSTPLNLPRISSEACEVNGYYVPTNTRLSVNIWAIGRDPDVWENPLEFNPE      420
VcF3'5'H      LCKEAFRLHPSTPLNLPRISSEACEVNGYYVPKNTRLSVNIWAIGRDPDVWENPLEFNPE      420
              ********************************.***************************

VmF3'5'H      RFMSGKNAKMDPRGNDFELIPFGAGRRICAGARMGVVMVEYFLGTLVHSFDWKLPDGVAE      480
VuF3'5'H      RFMSGKNAKMDPRGNDFELIPFGAGRRICAGARMGVVMVEYFLGTLVHSFDWKLPDGVAE      480
VcF3'5'H      RFMSGKNAKMDPRGNDFELIPFGAGRRICAGARMGVVMVEYFLGTLVHSFDWKLPDGVAE      480
              ************************************************************

VmF3'5'H      LNMDETFGLALQKAVPLAAKVTPRLHPSAYTM  512
VuF3'5'H      LNMDETFGLALQKAVPLDAKVTPRLHPSAYAM  512
VcF3'5'H      LNMDETFGLALQKAVPLAAKVTPRLHPSAYAM  512
              ***************** ************:*
```
