## Supplementary material for "Identification, characterisation and recombinant expression of flavonoid 3’,5’-hydroxylases and cytochrome P450 reductases from *Vaccinium* species": Suppl HPLC

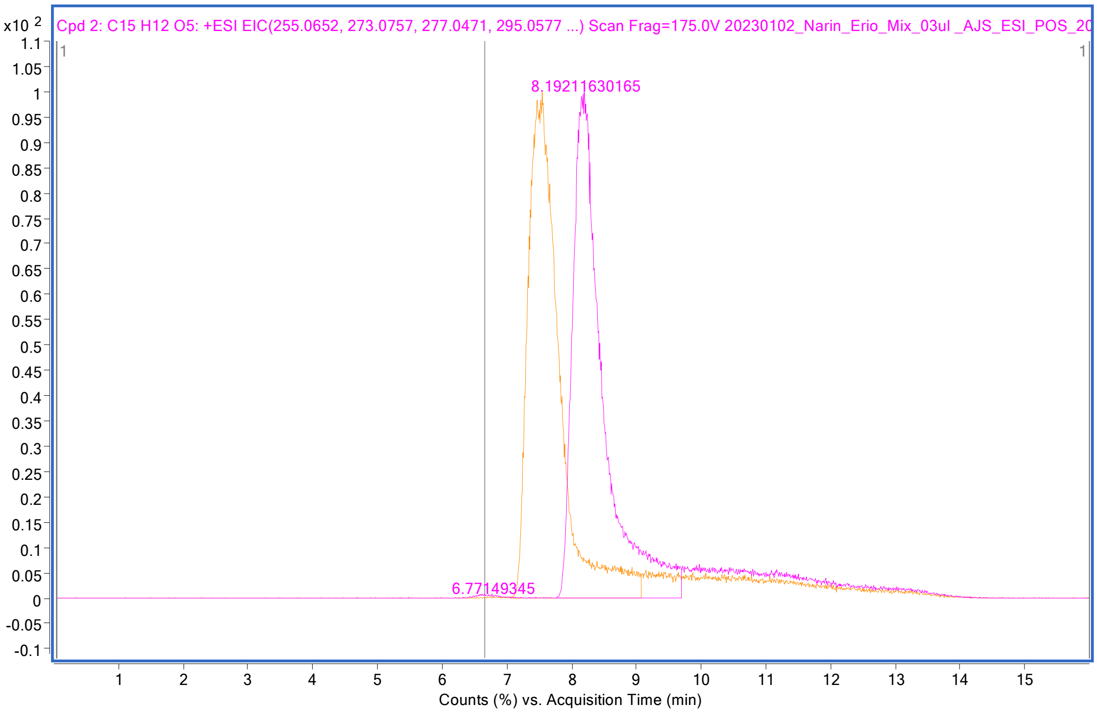


Supplementary figure 1. Elution profile of naringenin (pink peak)

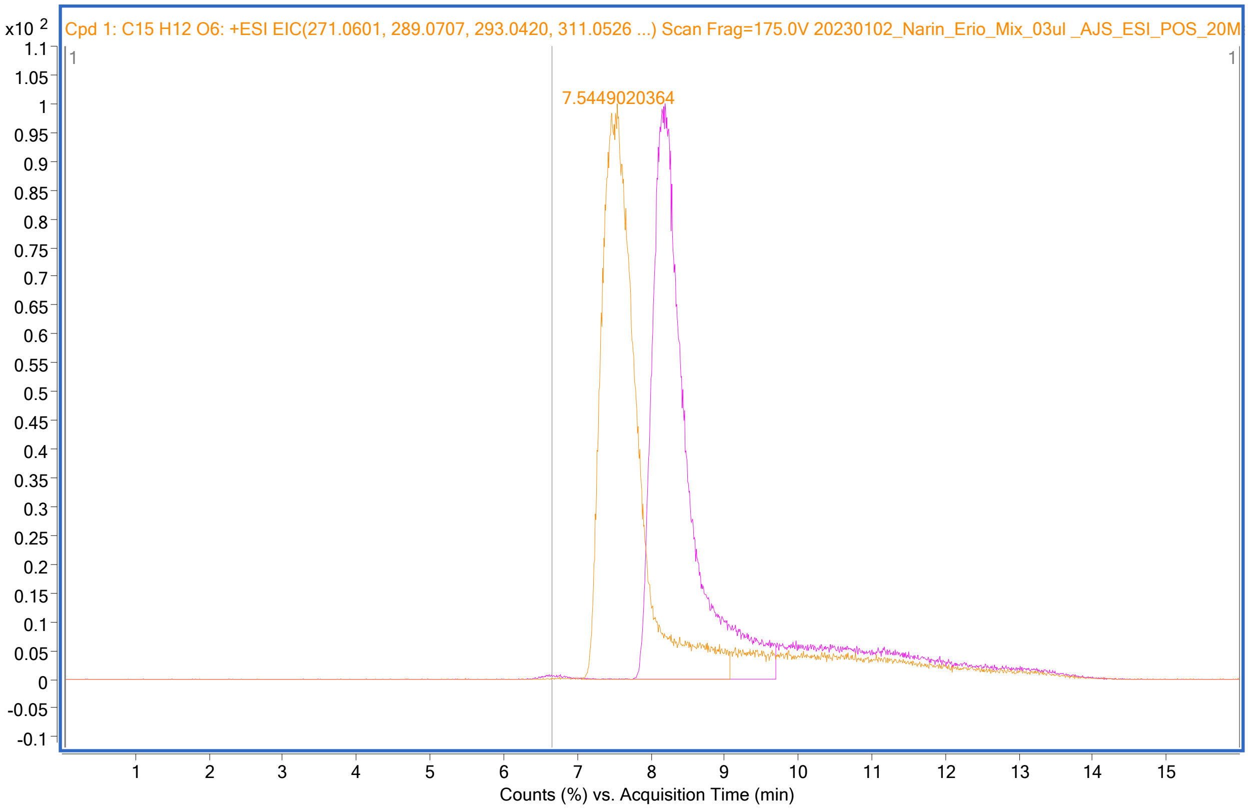


Supplementary figure 2. Elution profile of eriodictyol (orange peak)

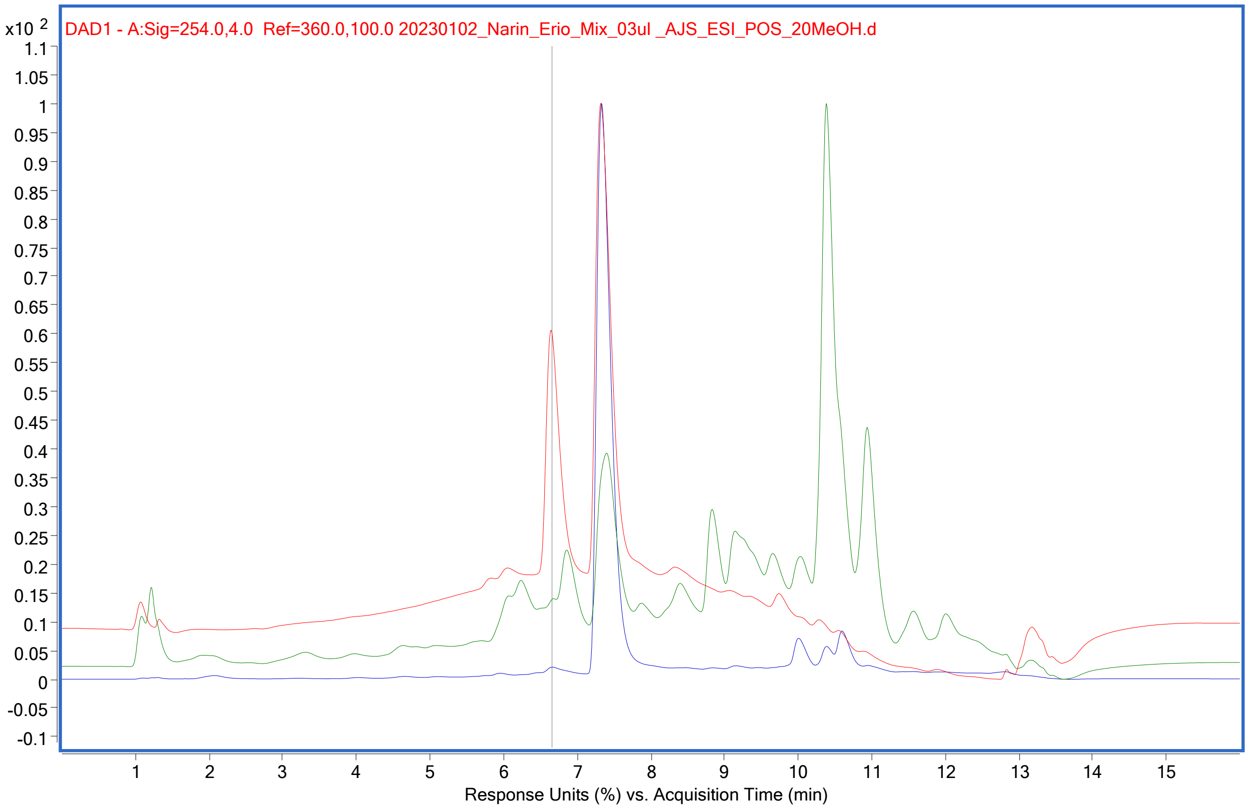


Supplementary Figure 3. Overlaid elution profiles showing that after incubating VuF3'5'H with VmCPR1, 10 µM naringenin and 1 mM NADPH only naringenin is detected. The red line corresponds to the standard mix of naringenin and eriodictyol, the green line shows the elution profile of the ethyl acetate extract from the enzyme assay detected at 254 nm, and the blue line at 290 nm.
