## Supplementary material for "Identification, characterisation and recombinant expression of flavonoid 3’,5’-hydroxylases and cytochrome P450 reductases from *Vaccinium* species": Suppl Table 1

SUPPLEMENTARY INFORMATION
**Table 1, Primers**

| Primer name | Primer sequence | Remarks |
|---|---|---|
| F35Hdegsense | ATGGCNAAYATGATHGGNCARGT | Degenerate primers used for amplifying fragment of F3′5′H |
| F35Hdegantis | GTRTCDGTDCCWGCDGTRAAYA | |
| 3-RACE-OligodT | AAGCAGTGGTATCAACGCAGAGTACTTTTTTTTTTTTTTTTTTTTTTTTTTTTTVN | 3'-RACE primers |
| 3-RACE-spec | AAGCAGTGGTATCAACGCAGAGT | |
| HaCPR5p | ATGCAATCAAGCTCCGAGAAGATC | Primers used for amplifying HaCPR |
| HaCPR3p | TTACCAAACATCACGGAGGTATCTTCC | |
| *Vm*CPR15p | ATGCAATCGAGCTCGGAAAAAGTG | Primers used for amplifying potential *Vm*CPR isoform 1 |
| *Vm*CPR13p | TCACCACACGTCACGCAAATAC | |
| *Vm*CPR25p | ATGGAATCGAGCTCCGTGAAGG | Primers used for amplifying potential *Vm*CPR isoform 2 |
| *Vm*CPR23p | CTACCAAACATCACGCAAATACCTC | |
| *Vc*F35HApaIKozakNtH6 | CAACTGGGCCCAAACGATGCACCATCACCATC | For amplifying *Vc*F3′5′H with an N-terminal 6xHis tag |
| *Vc*F35HCtermApaI | AGTATGGGCCCTTACATAGCATAAGCAGAAGGATGC | |
| *Vc*F35HApaIKozwoH6 | ACACTGGGCCCAAACGATGGCTCCAGATATTATGTTG | For amplifying *Vc*F3′5′H without an N-terminal 6xHis tag and stop codon |
| *Vc*F35HCtermnostopApaI | AGCAAGGGCCCCATAGCATAAGCAGAAGGATGCAA | |
| *Vm*F35HEcoRIKozNtH6 | GACATGAATTCAAACGATGCACCATCACCATCACCATGCCCTAGACATAATCTTGCTAAGG | For amplifying *Vm*F3′5′H with an N-terminal 6xHis tag |
| *Vm*F35HCtermEcoRI | ACGAGGAATTCTACATAGTATAAGCACTTGGATGCAG | |
| *Vm*F35HEcoRIKozwoH6 | ACTGTGAATTCAAACGATGGCCCTAGACATAATCTTGCTAAG | For amplifying *Vm*F3′5′H without an N-terminal 6xHis tag and stop codon |
| *Vm*F35HCtermnostopEcoRI | AGCTAGAATTCCATAGTATAAGCACTTGGATGCAGTC | |
| *Vu*F35HEcoRIKozNtH6 | GATACGAATTCAAACGATGCACCATCACCATCACCATGCCCTAGACATAATGTTGTTCAGG | For amplifying *Vu*F3′5′H with an N-terminal 6xHis tag |
| *Vu*F35HCtermEcoRI | GATACGAATTCTACATAGCATAAGCACTTGGATGC | |
| *Vu*F35HEcoRIKozwoH6 | GATACGAATTCAAACGATGGCCCTAGACATAATGTTGTTC | For amplifying *Vu*F3′5′H without an N-terminal 6xHis tag and stop codon |
| *Vu*F35HCtermnostopEcoRI | GATACGAATTCCATAGCATAAGCACTTGGATGCAG | |
| *Vc*F35HApaIwoTMH | ACACTGGGCCCAAACGATGTTGTTGAAGCCAACTTGTAAGTTG | For removing residues Ala2-Gln30 from *Vc*F3′5′H |
| *Vu*F35HApaIwoTMH | ACACTGGGCCCAAACGATGCTCCTTAAACCCACCTGTAAACTC | For removing residues Ala2-Gln30 from *Vu*F3′5′H |
| *Vu*F35HApaInostoprev | AGCTAGGGCCCCATAGCATAAGCACTTGGATGCAG | |
| *Vm*F35HApaIwoTMH | ACACTGGGCCCAAACGATGCTCCTCAAACCTGCCAGTAAACTC | For removing residues Ala2-Gln30 from *Vm*F3′5′H |
| *Vm*F35HApaInostoprev | AGCTAGGGCCCCATAGTATAAGCACTTGGATGCAGTC | |
| prAOX1MCSHis6TT FW | TGGACTGGTCTCGTGAGATCTAACATCCAAAGACGA | For amplifying the |

| | | |
|---|---|---|
| | AAGG | sequence from AOX1 promoter to transcription termination sequence of pPICZ A vector |
| prAOX1MCSHis6TT REV | TGGACTGGTCTCGGTGCACAAACGAACTTCTCACTTAATCTTCTGTACTCTGAAG | |
| pHygRt FWD | TGGACTGGTCTCCGCACTATATGTGAAGGCATGGCTATG | For amplifying hygromycin B resistance marker from pUDP082 vector |
| pHygRt REV | TGGACTGGTCTCCACCGTTGTTGAACATTCTTAGGCTG | |
| ori pAmpRt FWD | TGGACTGGTCTCCCGGTGAGCAAAAGGCCAGCA | For amplifying ampicillin resistance marker from pJET1.2 vector |
| ori pAmpRt REV | TGGACTGGTCTCCCTCAGGTGGCACTTTTCGGG | |
| VmCPR1ApaIfw | ACACTGGGCCCAAACGATGCAATCGAGCTCGGAAAAAG | For amplifying *Vm*CPR1 without an N-terminal 6xHis tag and stop codon |
| VmCPR1ApaIrev | AGCTAGGGCCCCCACACGTCACGCAAATAC | |
| VmCPR2ApaIfw | ACACTGGGCCCAAACGATGGAATCGAGCTCCGTG | For amplifying *Vm*CPR2 without an N-terminal 6xHis tag and stop codon |
| VmCPR2ApaIrev | AGCTAGGGCCCCCAAACATCACGCAAATACCTC | |
| HaCPRNotIfw | ATACTGCGGCCGCAAACGATGCAATCAAGCTCCGAGAAG | For amplifying *Ha*CPR without an N-terminal 6xHis tag and stop codon |
| HaCPRNotIrev | ATACTGCGGCCGCCCCAAACATCACGGAGGTATCTTC | |
| VmCPR1ApaINtH6fw | ACACTGGGCCCAAACGATGCACCACCACCACCACCACCAATCGAGCTCGGAAAAAGTG | For amplifying *Vm*CPR1 with an N-terminal 6xHis tag |
| VmCPR1ApaIstoprev | AGCTAGGGCCCTCACCACACGTCACGCAAATAC | |
| VmCPR2ApaINtH6fw | ACACTGGGCCCAAACGATGCACCACCACCACCACCACGAATCGAGCTCCGTGAAGGTG | For amplifying *Vm*CPR2 with an N-terminal 6xHis tag |
| VmCPR2ApaIstoprev | AGCTAGGGCCCCTACCAAACATCACGCAAATACCTC | |
| HaCPRNotINtH6fw | ATACTGCGGCCGCAAACGATGCACCACCACCACCACCACCAATCAAGCTCCGAGAAGATCTC | For amplifying *Ha*CPR with an N-terminal 6xHis tag |
| HaCPRNotIstoprev | ATACTGCGGCCGCTTACCAAACATCACGGAGGTATCTTC | |
| 5′AOX1sequencing primer | GACTGGTTCCAATTGACAAGC | Sequencing primers for yeast vectors |
| 3′AOX1sequencing primer | GCAAATGGCATTCTGACATCC | |
